# Phylogenetic signal in herbicide resistance evolvability, and the use of phylogenies to predict herbicide resistance risk

**DOI:** 10.64898/2026.04.23.720359

**Authors:** Neil Brocklehurst, Chun Liu

## Abstract

The evolution of herbicide resistance in weeds is a problem affecting both food production and ecosystems. Numerous factors affect selection towards herbicide resistance making prediction of resistance evolution complicated. The potential for a species to evolve resistance is linked to reproductive and metabolic traits, which may give resistance evolvability a phylogenetic signal and allow phylogenies to be used to predict resistance risk. Here we show that proxies for “resistance evolvability” show correlations with phenotypic traits that evolve at the deep-time level, including an aquatic lifestyle and pollination mode. This results in these proxies having a significant phylogenetic signal, demonstrating a deep-time evolutionary signal at the clade level is present. Machine learning models using phylogenetic distances as predictor variables are able to predict resistance risk with a high accuracy. Fitting macroevolutionary models based on Ornstein Uhlenbeck processes to the resistance evolvability proxies identifies four resistance regimes, including a separation between monocots and eudicots, with the latter evolving resistance less rapidly than the former. These analyses demonstrate the value of phylogenetic comparative methods in predicting resistance risk in weed species that become more problematic, and such methods may provide an invaluable addition to the numerous efforts to predict resistance evolution.

## Introduction

Weed management and control are vital in maintaining yields in agriculture, and herbicides provide an effective means of reducing weed populations. However, their use selects for resistant phenotypes in weed populations, creating both economic and ecological problems as resistant weeds harm food production and ecosystems (Powles & Yu 2010, Mortensen et al 2012). Successful assessment of the risk of resistance evolution, and prediction of the situations in which it is likely to arise, would therefore be invaluable.

The phylogenetic relationships between species provide a tool for predicting the evolution of traits (Baum & Smith 2013). The shared evolutionary history of species leads to greater phenotypic similarity between more closely related species (Darwin 1859; Harvey & Pagel 1991; Blomberg & Garland 2002). Traits that are more similar in more closely related species than in species drawn at random are said to show “phylogenetic signal”, a property that indicates the power of phylogeny in making predictions about that trait’s evolution (Blomberg & Garland 2002).

Herbicide resistance in weeds, along with resistance to other pesticides in pest species and antibiotics in bacteria, is a trait that evolves within recent populations, not in deep time at the clade level. Nevertheless, it is possible that the tendency for a species to evolve resistance (resistance “evolvability”) does show phylogenetic signal. The potential to evolve resistance is linked to various traits related to reproductive rate and mode (e.g. Jasieniuk et al 1996; Holt et al 2013; Tranel 2021), metabolic features (e.g. Yu & Powles 2014), and the evolutionary history of the genes associated with resistance (e.g. Christoffers 1999; Wang et al 2022).

Such traits have a deep time evolutionary history, and this will be reflected in their phylogenetic signal. If traits exhibiting phylogenetic signal are influential in increasing resistance evolvability, then resistance evolvability itself may show phylogenetic signal. Thus, if tendency to evolve resistance is found to show phylogenetic signal, then plant phylogeny may provide the basis for a predictive tool for assessing resistance risk.

Phylogenetic signal has been widely used in studies of ecotoxicology, showing phylogenetic signal in the susceptibility of various groups to pollutants, and the variation of the signal in different pollutants (e.g. Eriksson et al 2009; Jeffree et al 2010; Hammond et al 2012). It has also been incorporated into analyses of antibiotic resistance, demonstrating that the genetic material for antibiotic resistance has a macroevolutionary history dating from before the era of widespread antibiotic use (Aminov & Mackie 2007). Its study in connection with herbicide resistance has been limited, however, to analyses at higher taxonomic levels (Hill 1982, Holt et al 2013). Holt et al. (2013), for example, identified families where instances of resistance were overrepresented relative to others, finding such overrepresentation in Amararanthaceae, Poaceae and Brassicaceae. It should be noted, however, that this study represents an analysis treating phylogenetic relationships as a set of discrete Linnaean taxonomic units rather than as a continuous hierarchical set of relationships, with species separated by varying amounts of time. It also limits the potential for using the phylogenetic relationships in prediction of resistance risk to a coarse taxonomic level.

Here we use a time-calibrated species-level phylogeny to assess the strength of phylogenetic signal in herbicide resistance evolution in weeds. We examine the potential driving forces behind this signal, by demonstrating the correlation between resistance evolvability in species and various reproductive, life history and habit traits. We also demonstrate the potential for plant phylogenies to act as predictive tools in identifying clades where resistance is more or less likely to evolve. Finally, we make an exploratory effort at identifying resistance “regimes”: sections of the evolutionary tree united by similar resistance evolvability profiles.

## Materials and Methods

### Datasets

Recorded cases of resistance were downloaded in September 2022 from the International herbicide-resistant weed database (Heap 2022; http://www.weedscience.org). The dataset contains 1645 cases of resistance in 268 species across 72 countries and dates back to 1950s.

Lists of common and troublesome weed species in the USA and Canada were obtained from the WSSA surveys (https://wssa.net/wssa/weed/surveys/). These surveys ask public and private weed scientists, extension agents, crop consultants and other land and water managers with expertise in weed management to list common and troublesome weed species. The surveys are undertaken in three-year cycles, with different surveys carried out in consecutive years: first in broadleaf crops, fruit and vegetables, second in grass crops, and third in aquatic and non-crop areas. The cycle from 2019-2021 was used as the basis for this dataset (Van Wychen 2019, 2020, 2021), and the list of troublesome and common species were extracted. Taxa named to the genus level or higher were ignored, leaving 564 species.

### Phylogenetic framework

The 715 species from both datasets were organised into a timescaled phylogeny using an informal supertree approach to maximise taxon inclusion. The timescaled phylogeny of Li et al (2018) was used as a backbone constraint, being a recent analysis of angiosperm relationships and divergence time across a broad and comprehensive sampling of clades. Taxa that were not included in this backbone tree were added principally using data from the timetree 5 database (Kumar et al 2022; http://www.timetree.org/), which contains data on relationships and divergence times of species from across the tree of life. A search requesting the divergence date between two species will produce a list of molecular clock studies that have included either the species themselves or representatives of the higher taxa to which they belong. These dates can be used to infer the relationships between species and a divergence date of each node in the tree may be selected from those given.

Where information on relationships and divergence dates of a species was not obtainable from Timetree or by a survey of the published literature, or where the only dates available are inconsistent with those of Li et al. (2018), then the relationships from published trees were used but the relevant nodes in the tree were not assigned a date (the date to be inferred below). Where no information on the phylogenetic relationships between certain species were available, they were left as an unresolved polytomy (to be resolved below). The phylogeny in nexus format, the ages assigned to each node, and the justification for the node ages and relationships used, are presented in supplementary data 1 and 3.

Having generated a phylogeny and list of ages for the internal nodes, the unresolved relationships and undated nodes were inferred using the Birth Death Skyline model (Stadler et al 2013). This is a Bayesian approach that simultaneously makes estimates of speciation and extinction rates and the relationships and node dates. This was carried out in MrBayes V3.2.7 (Romquist et al. 2012). The relationships were constrained to the inferred tree and the node ages were fixed at those defined above. The analysis consisted of four runs with four chains with 10000000 generations, sampling every 1000. The maximum clade credibility (MCC) tree was used for subsequent analyses. The analysis file is presented in supplementary data 2 and the maximum clade credibility tree in supplementary data 3.

### Traits

Six traits were selected to test for their correlation with resistance evolvability, representing various aspects of the plants’ reproduction, growth habit, lifestyle, and genetics. These six traits are not meant to be an exhaustive list of the traits that could impact a plant’s resistance evolvability but were selected as varied examples of traits for which data was available for the majority of species in the dataset (Supplementary Data 13).

Pollination mode indicates the means by which the plant may spread the genes for resistance within its population or to other populations, and the likely distance over which resistance may spread (Tranel 2021; Ambruster 2023). Self-compatibility also affects the spread of the resistance genes; self-compatible plants are more likely to produce offspring with the resistant genotype (Gressel 2009). However, self-incompatibility, forcing outcrossing, means it is more likely the resistant genes may spread to other populations; the latter point has been noted as a potential reason why herbicide resistant populations of *Amaranthus tuberculatus* are able to spread so rapidly (Tranel 2020). Pollination mode was assessed as a discrete trait (Animal, wind and self), while self-compatibility was treated as binary (compatible or incompatible).

Reproductive longevity (annual, biennial or perennial) has been found to impact frequency of resistance evolution, with annual weeds being more likely to evolve resistance (Holt et al. 2013). The precise reasons are unclear; Holt et al. (2013) attributed it to species with shorter lifespans reacting more rapidly to directional selection. However, this factor should only apply to shorter reproductive lifespans, and the majority of weedy perennials (which are mostly herbaceous rather than woody shrubs or subshrubs) are able to reproduce in their first season. An alternative explanation is that annuality is correlated with a life-history strategy that trades off long-term survival and growth with maximising reproductive potential in one season e.g. higher seed counts, greater frequency of self-fertilisation (Stebbins 1950; Pitelka 1977; Primack 1979; Vico et al. 2016; Barrett & Harder 2017). This in turn leads to greater rates of evolution and more rapid spread of the resistance genes (Andreason & Baldwin 2001; Smith & Donoghue 2008). Reproductive longevity was assessed as a discrete trait: Annual, Biennial and Perennial.

The relationship between genome size and rates of evolution is complicated and not fully understood, but it is considered that larger genomes can lead to greater rates of evolution (Van de Peer et al. 2007; Freeling & Thomas 2006). Genome size here was represented by the 2C value downloaded from the plant DNA C values database (Leitch et al. 2019; https://cvalues.science.kew.org/), with values treated as continuous.

Growth habit was included as a trait that has been linked to “weediness”, if not resistance evolution, with fewer weeds being shrubs and more being forbs and graminoids (Kuester et al 2018). Woody shrubs are likely less weedy due to their slower growth and reproduction, a factor that should also affect their tendency to evolve resistance. Also, if fewer shrubs are weedy, they are less likely to be exposed to herbicides. Four growth habits were considered: Herbaceous/forb, Graminoid, vine, subshrub and shrub.

An aquatic lifestyle was also noted by Holt et al. (2013) as being linked to less frequent resistance evolution. This was not thought to be due to a trait or feature of aquatic plants that might affect rates of evolution or reproductive strategies. Instead, it was thought to be due to the less frequent use of herbicides in aquatic systems limiting the selection pressure towards resistance (Rasodovich et al 2007; Holt et al. 2013). Nevertheless, even though the cause of the less frequent resistance in aquatic species is thought to be due to agronomic practices rather than some intrinsic factor, this does not preclude this trait from consideration in the context of phylogenetic signal of resistance evolvability; it is still to be expected that an aquatic lifestyle is a trait with phylogenetic signal, and this would still impart phylogenetic signal to resistance evolvability.

### Proxies for Resistance Evolvability

Three proxies were used to quantify a species’ tendency to evolve resistance. These were intended to measure the rapidity and frequency of resistance evolution, while accounting for the fact that certain species will, due to their abundance, geographic range, and range of situations in which they are competitive (aspects of “weediness” rather than tendency to evolve resistance), be exposed to more herbicidal active ingredients and modes of action more frequently, potentially driving increased frequency of resistance evolution independent of resistance “evolvability”. There is also the fact that the frequency and intensity of resistance screening is inconsistent across countries, being most frequent in areas where there are more active scientists with the resources to study resistance (Peterson et al 2018). The quality of sampling (the frequency and intensity of screening for resistance) potentially impacts the relative frequencies of resistance observations. This can lead to uncertainty regarding how genuine the frequencies patterns observed are.

The first proxy represents a count of resistance cases in each species but subsampling each country to a consistent coverage to account for variation in sampling. Subsampling to fixed levels of coverage has been shown to account more accurately for sampling heterogeneity than sampling to fixed sample sizes (Alroy 2010; Chao & Jost 2012). Coverage of the species underlying frequency distribution is estimated using Good’s U (Good 1953).

Subsampled resistance counts were calculated for each species by adapting the algorithm of Alroy (2014). For each country, all resistance cases recorded from that country by the International Herbicide-Resistant Weed Database (data downloaded September 2022, Supplementary Data 5) were drawn sequentially at random, tracking the value of Good’s U. Each time the target coverage was crossed, the number of cases drawn from each species were counted. After all cases were drawn, the mean cases from each species were calculated. The subsampled cases from each species counted from each country was summed to produce a global count of resistance cases for each species. This approach was repeated 1000 times and the mean count was used to represent the proxy for resistance frequency. This was carried out using custom code written in R v4.1.3 (R Core Team 2022), presented in Supplementary Data 6.

The second proxy was an attempt to quantify the rapidity with which a species tends to evolve resistance, regardless of the frequency with which it is targeted by herbicides and the variety of herbicides to which it is exposed. For each species, the year of first appearance of resistance to an active ingredient was subtracted from the year of introduction of that ingredient. A median time to resistance was then calculated for all actives to which that species had recorded resistance to. The intention behind this proxy was to provide an estimate of the rapidity with which a species tends to evolve resistance on the occasions where it is exposed to herbicides, however infrequently. Years of introduction of the active ingredients were drawn from Pesticide Properties DataBase, developed by the Agriculture & Environment Research Unit at the University of Hertfordshire (http://sitem.herts.ac.uk/aeru/ppdb/en/index.htm; Supplementary data 7).

The third proxy was based on the WSSA survey data. The weeds identified as common and troublesome in the USA and Canada are those most likely to be subjected to frequent herbicide use and therefore experience the selection pressures towards evolution of herbicide resistance. Therefore, those which are recorded as having evolved resistance in these countries may be assumed to represent those with greater resistance evolvability. The binary trait of resistance recorded/not recorded in the USA or Canada was used as a resistance proxy within species recorded in the WSSA survey only (Supplementary data 8).

### Correlation between traits and resistance evolvability

The phylogenetic generalised least squares method (Grafen 1989; PGLS) was used to test the correlation between the six traits and the two continuous resistance proxies. PGLS is a modification of generalised least squares analysis that incorporates the phylogenetic relationships of taxa as an expected covariance. PGLS allows both single variables and multi-variate models to be compared to the dependent variables, while not violating the assumption of independence of data points due to the phylogenetic non-independence of species.

In this case models incorporating one, two and three predictive variables, with all possible variable combinations, were tested. Models containing greater number of variables were not tested due to the number of species in the dataset; phylogenetic comparative methods are unreliable when fewer than 40 taxa per free parameter are available (Freckleton et al 2002). The model fits were compared using the Akaike weights. For the traits that appear in models with greater support than the Null, values of the resistance proxies for plants appearing in each category were compared using Mann Whitney U tests.

### Analyses of Phylogenetic Signal

Two metrics were used to calculate the phylogenetic signal of the resistance evolvability proxies. For the two continuous proxies, Pagel’s λ (Pagel 1999) was employed, due to simulations demonstrating it performing better than other metrics particularly under more complex models of evolution (Münkemüller et al 2012). It was implemented using the *phylosig()* function in the R package phytools (Revell 2012). The proxy values were log transformed prior to analysis. The significance of the phylogenetic signal was assessed using a likelihood ratio test comparing the fit of the lambda model (where internal branches are scaled by the lambda value) to one where lambda is 0 (no phylogenetic signal).

For the discrete proxy, Fritz & Purvis’s D (Fritz & Purvis 2010), a measure of phylogenetic signal in discrete binary traits, was used, implemented in the R package caper (Orme et al 2013). Significance was measured by comparisons with a null distribution, where D values are calculated for a trait set drawn at random from a normal distribution.

### Prediction of Resistance using Phylogeny

There are numerous methods to predict values of evolutionary traits from phylogenies. Multiple approaches were tested, each validated using K-fold cross validation: the dataset was divided into 10 folds at random; each fold was, one by one, used as a pseudo-test dataset, with data from the remaining folds used to predict values of the resistance proxy in the pseudo-test dataset. This produces a predicted value of the resistance proxy for each data point, which may be compared to the observed values. For the two continuous proxies, the predicted values were compared to observed values using the Pearson’s product moment coefficient, and for the discrete proxy a confusion matrix was created.

Four approaches were tested for predicting the values of the continuous proxies: three machine learning regression approaches and a likelihood model fitting approach. The machine learning approaches used the patristic distances between a taxon and all taxa in the training dataset as predictors for the test variable (the resistance proxy). Under a Brownian Motion model of evolution, the amount of change expected along a branch is directly proportional to the length of the branch in time (Rohlf 2001; Revell et al. 2008). Therefore, in a phylogeny scaled to time, the patristic distance between two taxa (the sum of the lengths of the branches separating them) may act as a predictor for the amount of trait variation between them. Machine learning models linking these distances to the resistance proxies were built using three methods: Random Forest, Generalised Linear Models, and Support Vector Machines, with the models build and tuned using the caret package in R. The fourth approach was to use the maximum likelihood model fitting approaches commonly employed in ancestral reconstructions of continuous traits. A Brownian Motion model is fit to the observed trait data and phylogeny, and used to infer the expected median and variance of values of that trait at the internal nodes (Felsenstein 1973, Schluter et al 1997). The *anc.ML()* function in phytools allows this approach to be used to infer not only the trait values at the internal nodes but also at tip nodes where the trait values are missing.

For predicting the discrete proxy, the same three machine learning approaches were tested, along with two others. The first is the SIMMAP approach of stochastically mapping a discrete trait’s evolution over a phylogeny (Bollback 2006). Maximum likelihood is used to fit a character state transition model to the phylogeny, and this model and the tip states are used to simulate evolutionary histories of the character over the tree. For each fold, 1000 simulations were generated with the pseudo-test taxa given equal prior probabilities of resistant/not resistant. The approach was implemented using the *make.simmap()* function in phytools. The final approach employed the threshold model. This model treats a discrete trait as a presentation of an underlying continuous trait called the “liability” which evolves under Brownian Motion. When the liability crosses a particular threshold, the discrete character state changes (Felsenstein 2012; Revell 2014). A Bayesian MCMC approach is used to estimate liabilities within the tree and thresholds between states, and thus infer character states for internal nodes and taxa for which data is missing (Revell 2014). This was implemented using the *ancThresh()* function in phytools. The analysis was run for 10,000,000 generations, sampling every 1000.

### Identification of Resistance Regimes

The Surface algorithm (Ingram & Mahler 2013) locates a set of macroevolutionary regimes characterised by Ornstein Uhlenbeck models of evolution, where the trait is drawn to a particular adaptive optimum (θ). The regimes identified by the analysis have different values of θ for each trait under investigation. The algorithm uses likelihood to fit a model with two regimes, identifying the best node within the tree to place the regime shift according to the Akaike information criterion (AIC). It then holds the position of that shift and iteratively searches for further shifts until the AIC values no longer improve. The algorithm then tests whether merging regimes improves the AIC score, indicating where regimes evolved convergently.

The algorithm was applied to both the continuous resistance proxies to identify resistance regimes, using functions in the R package surface (Ingram & Mahler 2013). The maximum number of shifts was constrained to 5 to avoid overfitting, following recommendations that, for phylogenetically non-independent data, one needs at least 40 taxa per free parameter to accurately estimate (Freckleton et al 2002).

### Sensitivity analyses

Estimates of phylogenetic signal may be influenced by variations in the evolutionary tree used in the analysis, including different relationships and estimates of node ages. To show how this may affect the estimates of phylogenetic signal, the analysis was repeated with alternative tree topologies and age estimates. First, 1000 trees were drawn at random from the posterior distribution produced by the birth death skyline analysis (Supplementary Data 4).

These trees will have the same backbone constraint as the MCC tree used in the analysis but will have different divergence time estimates for those nodes where constraints were not applied, and different relationships produced by the resolution of polytomies. To examine the impact of larger variations in node ages and relationships, a second supertree was created. This used the Timetree master-tree (Kumar et al 2022) as its backbone instead of the phylogeny of Li et al (2018). Each node in this master-tree was dated by finding the median of all relevant divergence time estimates in their database, with a smoothing technique applied to resolve node age conflicts Kumar et al (2022). The 190 taxa not found within the Timetree database were added to the tree based on published phylogenetic analyses, in the manner described above (Supplementary data 9). The tree was then subjected to the Fossilised Birth Death Skyline analysis (Supplementary Data 10, 11), with the same parameters described above, and 1000 trees were drawn from the posterior sample (Supplementary Data 12).

In total, therefore, 2000 trees representing a variety of possible evolutionary relationships and sets of divergence dates, were used in the sensitivity analyses. These were subjected to the analyses of phylogenetic signal to show how these parameters vary the results. The results are presented in Figure S1.

## Results

### Correlation between resistance evolvability and traits

No single model of those tested receives overwhelming support relative to others according to the Akaike weights. The single best fitting model receives an Akaike weights score on 0.15, and the combined Akaike weights score of all models with greater support than the null is only 0.35 (Table 1). The best fitting model correlates the resistance evolvability proxies with an aquatic lifestyle. The second is a multivariate model including both an aquatic lifestyle and pollination mode, and the third pollination mode only. The fourth is the null (Table 1).

**Table 1:** The model fit of the best fitting models identified by phylogenetic least squares, comparing the proxies for resistance evolvability to plant traits.

| Model | LnL | AICc | Akaike weights |
| --- | --- | --- | --- |
| Resistance Evolvability ~ Aquatic lifestyle | -150.05 | 304.18 | 0.15 |
| Resistance Evolvability ~ Aquatic lifestyle + Pollination mode | -148.07 | 304.43 | 0.13 |
| Resistance Evolvability ~ Pollination mode | -149.69 | 305.56 | 0.07 |
| Null | -151.95 | 305.93 | 0.06 |
| Resistance Evolvability ~ Aquatic lifestyle + Genome size | -150.04 | 306.26 | 0.05 |
| Resistance Evolvability ~ Aquatic lifestyle + Self compatibility | -150.04 | 306.26 | 0.05 |

According to a Mann Witney test, aquatic weeds have significantly fewer cases of resistance than non-aquatic weeds (Figure 1, Table 2). However, the aquatic weeds do have significantly shorter times to resistance appearing. The same test shows that wind and self-pollinated plants have significantly more frequent evolution of resistance than animal-pollinated plants, although not more rapid (Figure 1, Table 2).

**Figure 1:**
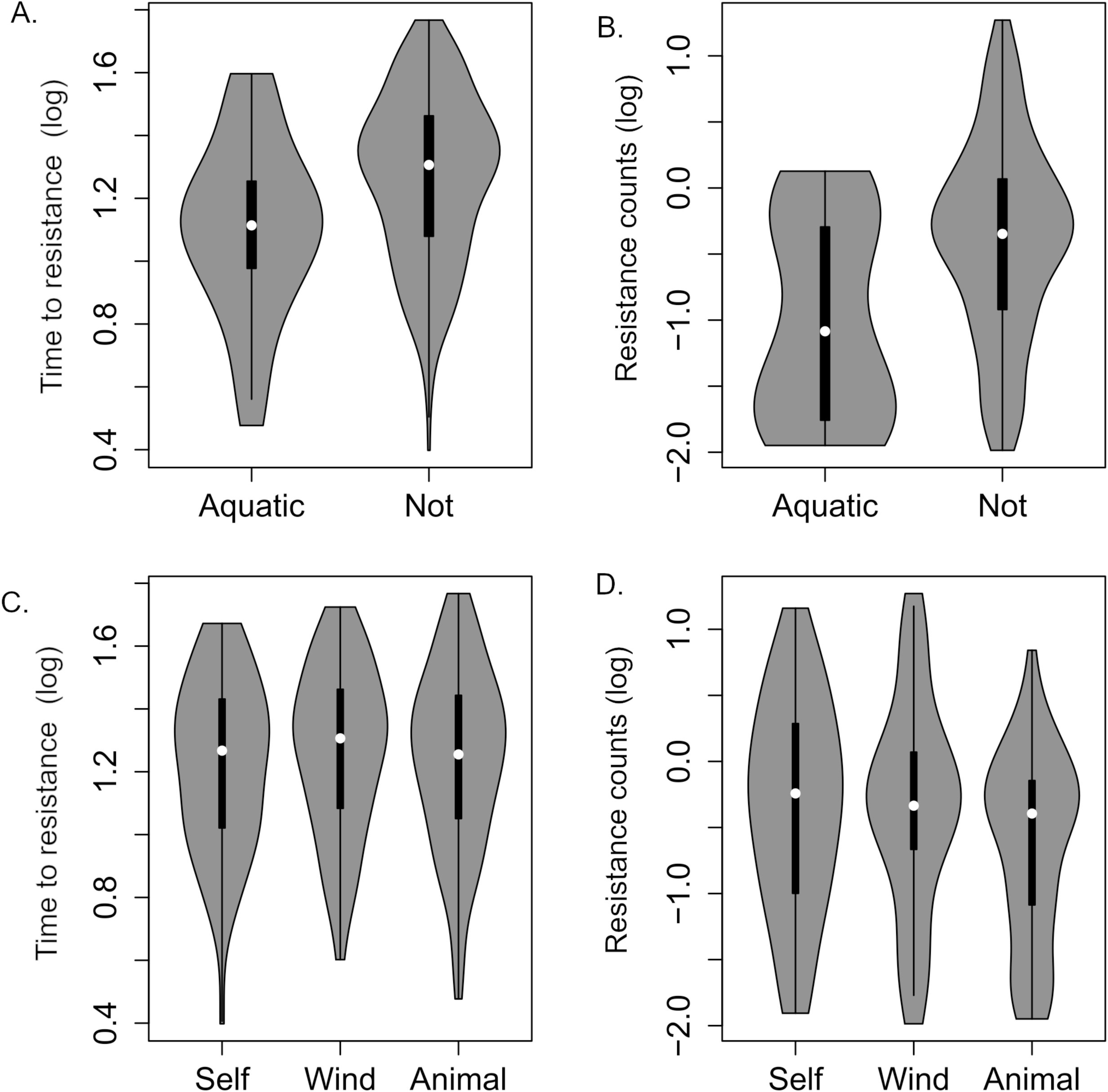
Violin plots comparing time to resistance (A,C) and resistance counts (B,D) to plants with aquatic/non-aquatic lifestyles (A,B), and different pollination modes (C,D)

**Table 2.** Results of the Mann Witney tests comparing time to resistance and resistance counts to plants with aquatic/non-aquatic lifestyles and different pollination modes.

| Resistance evolvability proxy | Trait comparison | U (p value) |
| --- | --- | --- |
| Time to resistance | Aquatic vs non-aquatic | 2321 (0.0027) |
| Resistance counts | Aquatic vs non-aquatic | 1172 (0.0003) |
| Time to resistance | Self-pollinated vs wind pollinated | 2653 (0.32) |
| Resistance counts | Self-pollinated vs wind pollinated | 2830 (0.92) |
| Time to resistance | Self-pollinated vs animal pollinated | 3075 (0.63) |
| Resistance counts | Self-pollinated vs animal pollinated | 3556 (0.024) |
| Time to resistance | Wind pollinated vs animal pollinated | 4651.5 (0.6) |
| Resistance counts | Wind pollinated vs animal pollinated | 4961 (0.02) |

### Phylogenetic signal in proxies for resistance evolvability

The phylogenetic signal of resistance counts and time to resistance was assessed by Pagel’s lambda as 0.37 and 0.41 respectively. These values, while not high, were significantly higher than would be expected were the traits distributed at random over the tree (Table 3). The relatively low values might be an indication that the tendency to evolve resistance is largely mediated by factors outside the scope of weed biology, such as agronomic practices and herbicide chemistry. Nevertheless, there is still a significant phylogenetic signal, indicating that biological factors such as reproduction and metabolism are significant factors in driving the frequency and speed of weed resistance (Figure 2).

**Figure 2:**
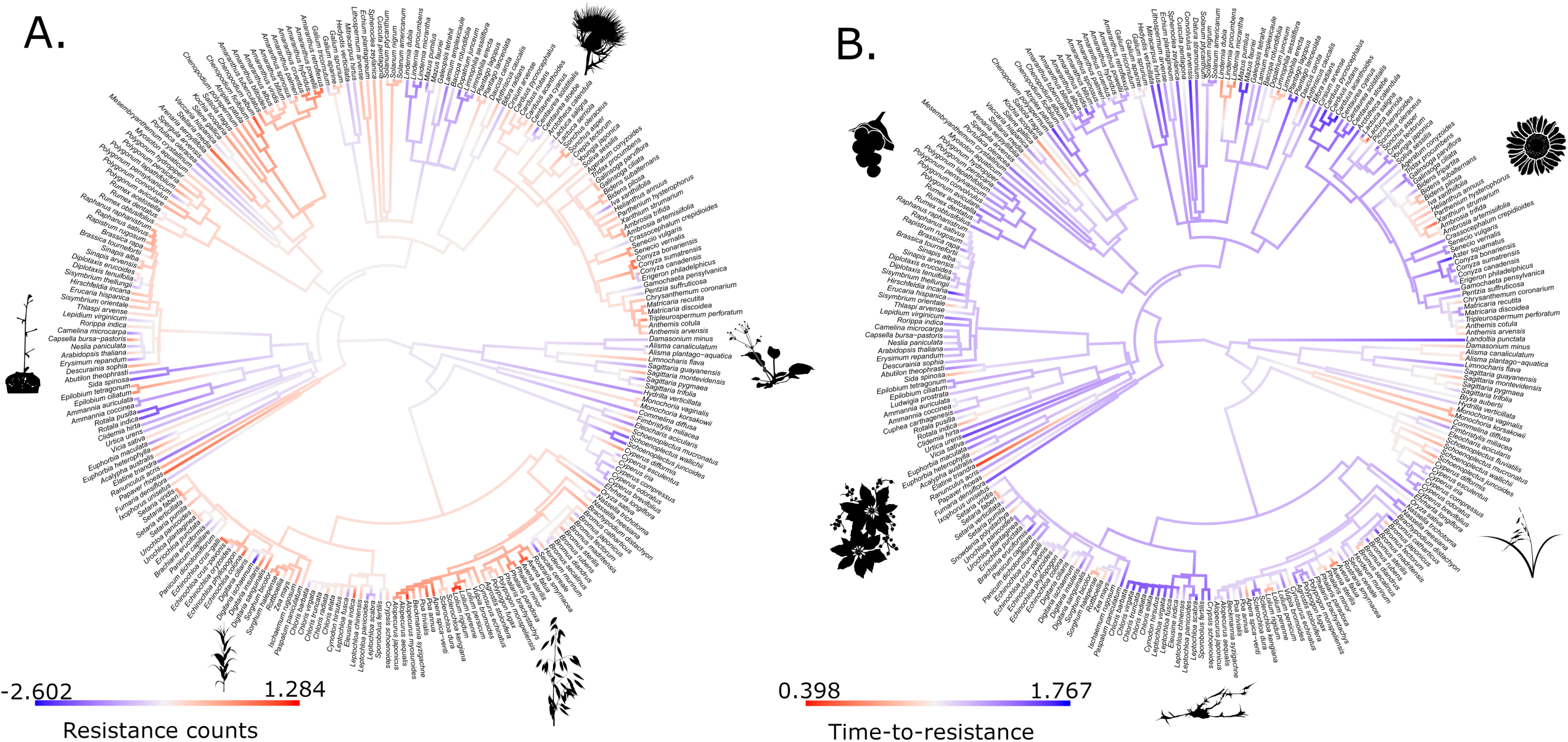
Resistance evolvability proxies mapped over the phylogeny. A) Resistance counts. B) Time-to-resistance. In both cases the red end of the colour scale indicates high resistance evolvability (high values of resistance counts but low values of time-to-resistance). Traits were mapped onto phylogeny using likelihood ancestral state reconstruction with the *contMap()* function in the R package phytools.

**Table 3:** Phylogenetic signal of the three resistance proxies. Pagel’s λ represents phylogenetic signal of continuous traits. Fritz & Purvis’s D represents phylogenetic signal of discrete binary traits.

| Proxy | Pagel's $\lambda$ | Fritz & Purvis's D | P value |
| --- | --- | --- | --- |
| Resistance Counts | 0.37 | NA | $1.61 \times 10^{-6}$ |
| Time-to-resistance | 0.41 | NA | 0.039 |
| Resistance in USA & Canada | NA | 0.69 | 0 |

Resistance evolution recorded in North America has a relatively higher phylogenetic signal, with a Fritz & Purvis’s D value of 0.69. As with the other proxies this is significantly higher than would be expected were the traits distributed at random with respect to phylogeny (Table 3).

### Prediction of Resistance Risk

Of all the methods tested to predict the resistance proxies from the phylogeny, K-fold cross validation consistently identified models built using the Support Vector Machine (SVM) approach as the most reliable predictors. SVM regression, using only the phylogenetic distances as predictor variables, was able to predict both resistance counts and time to resistance with a high degree of accuracy. The correlation between predicted values and true values, measured by the Pearson correlation coefficient, is 0.93 (p = < 2.2×10^-16^) for time to resistance and 0.92 (p = < 2.2×10^-16^) for resistance counts (Figure 3).

**Figure 3:**
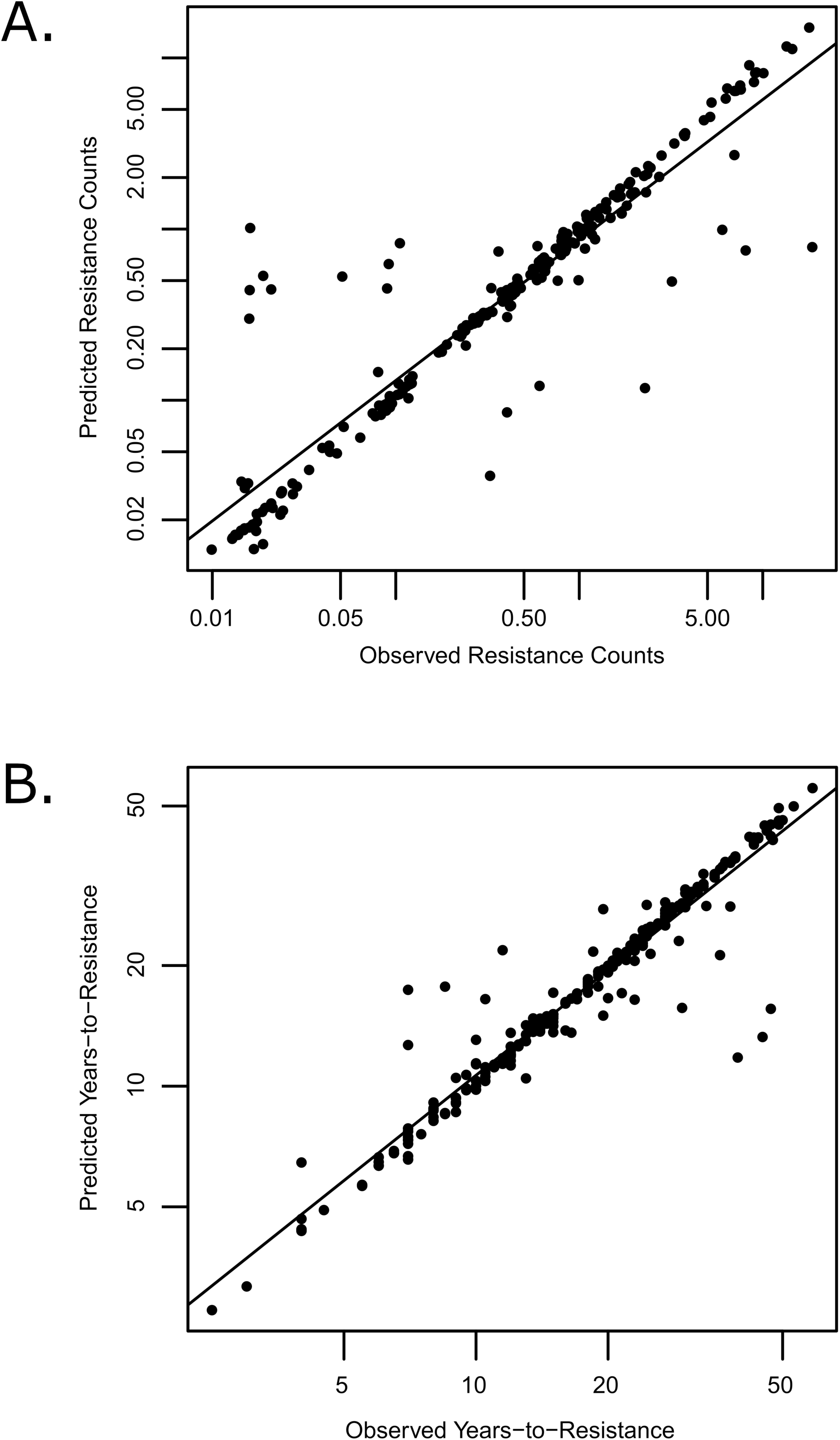
A comparison of values of resistance evolvability proxies observed in species to those predicted by SVM models during K-fold cross validation. A) Observed and predicted values of Resistance Counts; B) Observed and predicted values of years-to-resistance.

SVM classification of presence of resistance in America also shows reasonably accurate prediction of resistance and sensitivity. Of the 97 species from which resistance was recorded in the USA and Canada, 86 were predicted to show resistance (89% accuracy). Of the 467 common and troublesome weed species in which no resistance has been recorded in these countries, all were predicted to show no resistance (Table 4).

**Table 4:** Confusion matrix comparing the numbers of true and false predictions of resistance observations via the SVM model. Each cell contains the absolute number of species for which the observation and prediction have been made.

|  | No resistance predicted | Resistance predicted |
| --- | --- | --- |
| No resistance observed | 467 | 0 |
| Resistance observed | 11 | 86 |

### Identifying Resistance Regimes within plant phylogeny

Applying the Surface algorithm to resistance counts and time to resistance identifies four regimes (Figure 4). The values of θ for each proxy in each regime are presented in Figure 4B. Regime 1 represents the Eudicots. The shift to regime 2 occurs at the origin of the monocots. There is an increase in the frequency of resistance and a decrease in the time to resistance relative to regime 1. The shift to regime 3 occurs within regime 2, at the origin of the grass clade Chloridoideae. The frequency of resistance is similar to that of regime 1, but the time to resistance is longer. Two shifts to Regime 4 occur independently within regime 1: within Helianthae and the genus *Lindernia*. This regime has low values of both resistance counts and time to resistance.

**Figure 4:**
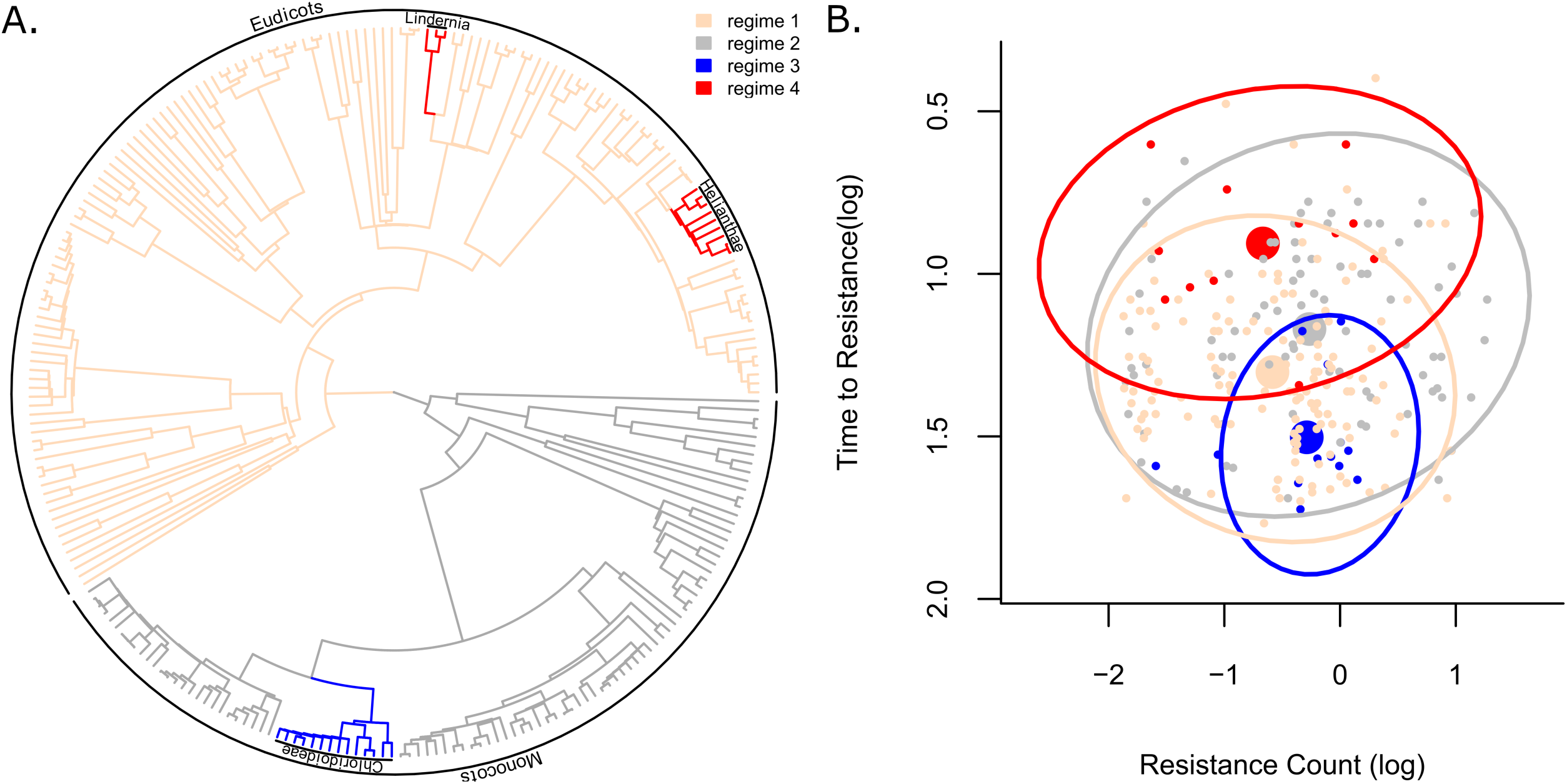
Resistance regimes identified by the surface algorithm. A) The four resistance regimes mapped onto plant phylogeny; B) The regimes mapped against trait values. Small points represent individual species. Large points represent the θ values (adaptive optima) of each regime. Ellipses represent 95% confidence ellipses.

## Discusssion

### Drivers of Resistance Evolvability

The two traits which PGLS identified as fitting the resistance evolvability proxies better than the Null model, whether on their own or as a multivariate model, are an aquatic lifestyle and pollination mode. The fact that wind-pollinated plants are more likely to evolve resistance than animal-pollinated plants is expected and has been noted before; wind pollinated plants experience wider gene flow, allowing resistance alleles to spread to novel populations, and also have greater reproductive success (Friedman & Barrett 2009; Hulme & Liu 2021).

The increased frequency of primarily self-pollinated plants over animal-pollinated plants is more unexpected; in the past obligate outcrossing has been associated with the increased spread of resistance (e.g. Jasienuik et al 1996; Tranel 2020; Hulme & Liu 2021). On the other hand, a distinction does need to be made between the spread of resistance spatially (i.e. where the resistant genes move greater distances) and the increase in abundance of the resistance genes at the local level. Species able to self-pollinate are more likely to have advantageous alleles increase in frequency, or achieve fixation, within the population more rapidly than outcrossing species (Kreiner et al. 2018). This is especially the case where the relevant trait is recessive or where there are differences in selective pressure between homozygotes and heterozygotes; increased selfing will increase the frequency of homozygosity, and the reduced gene flow will reduce the dilution of the resistance alleles.

The influence of an aquatic lifestyle on resistance evolvability is more difficult to explain. The aquatic weeds have significantly fewer cases of resistance than non-aquatic weeds, but evolve resistance significantly more rapidly. Holt et al. (2013) did note that aquatic weeds were under-represented in the IHRWD, and suggested that this was due to the reduced use of herbicides in aquatic systems. Almost all the cases of resistance in aquatic weeds occur in rice-growing situations, where the use of herbicides has historically been less prevalent and hand-weeding has been more widely used (Naylor 1994). Rice growers are also less varied in their herbicide use, the aquatic setting necessitating herbicides with low volatility and solubility, and the portfolio is dominated by Acetolactate Synthase inhibiting herbicides (ALS) (Strom 2018; Hamamura 2018). The more limited portfolio provides fewer opportunities for resistance to evolve, as they are exposed to fewer actives and modes of action.

However, while this does provide an explanation for the significantly fewer resistance cases observed in aquatic weeds, it does not explain the more rapid evolution of resistance in these weeds. This is also supported by the analysis of resistance regimes, with *Lindernia* fitting the regime of infrequent but very rapid resistance evolution *Lindernia* is a troublesome weed genus in rice paddies (Uchino & Watanabe 2002; Yoshino et al 2006; Kwon et al 2011), and it is only in this situation that resistance has been recorded in the IHRWD. The rapid evolution of resistance in aquatic settings may be due to the limited herbicide portfolio discussed above; with few options, the weeds will experience persistent exposure to a limited number of active ingredients. Herbicides may persist at low levels in water and aquatic sediments, providing continuous low-rate exposure to the weeds and gradually selecting for increased tolerance and resistance (Zanella et al. 2011, Muir 2024). There may also be influence of the prevalence of ALS herbicides used in rice growing, which is a mode of action that seems to have a high tendency to drive resistance evolution (Tranel & Wright 2002, Holt & Thill 1994; Blackshaw et al. 1994). Resistance to ALS-inhibiting herbicides was found in 1987, less than five years after their introduction (Mallory-Smith et al. 1990), and the frequency of cases has increased more rapidly than other widely-used herbicide groups (Tranel & Wright 2002), now represent the most frequent cases of resistance (676 cases in the IHRWD).

However, the potential for an explanation linked to the phenotype and ecology of aquatic plants should not be overlooked. Aquatic plants have substantially different metabolisms and reproductive strategies to terrestrial plants. Movement of chemicals via xylem and phloem is more limited in aquatic plants, and in many the vascular tissue is vestigial (Arber 1920; Ware et al. 2023, McDonald 2011). This could potentially affect the plant’s ability to evolve mechanisms of sequestering herbicides or limiting uptake. Aquatic plants frequently employ clonal reproduction, allowing rapid proliferation of resistant individuals (Barrett 1989, Eckert et al. 2016); as discussed above such reproduction can more rapidly allow the relevant alleles to reach fixation in a population. Their life-history traits may also be a contributing factor, as they often grow rapidly and show precocious reproduction, allowing shorter generation times and more rapid evolution (Barrett et al. 1993; Barrat-Segratain 1996).

### Prediction of Resistance Risk

The link between certain phenotypic, genetic and ecological traits and the tendency of a plant to evolve resistance obviously provides an avenue for prediction of resistance evolution, as has already been discussed in the past literature. However, this fact has a further implication; since these traits are those which have evolved in deep time, they, and therefore resistance evolvability itself, will have a phylogenetic signal, where more closely related species are more likely to share similar levels of resistance evolvability. The advantage of a phylogenetic classification system, based as it is on the evolutionary history of a clade, is that it allows explicit predictions to be made about the anatomy, physiology and genomics of individual species (Baum & Smith 2013). The fact that a species’ tendency to evolve resistance to herbicides has a significant phylogenetic signal demonstrates that phylogeny has predictive power for this trait. This potentially has immense value for weed control: as new weed species become more abundant or troublesome, their position in plant phylogeny may be used to identify how likely they are to show resistance to herbicides, allowing patterns of resistance across different plant lineages and factors influencing its evolution to be better studied, and allowing screening efforts to be more targeted.

The machine learning models created in this study, which predict resistance proxy values for species based on their phylogenetic distance from other species, are found by K-fold cross validation to show a high degree of accuracy in their predictions. The predicted values for the time-to-resistance proxy and the resistance frequency proxy correlate very accurately with those observed. The prediction of resistance cases in America is also accurate, with no false positives and a balanced accuracy (mean of the true positive and true negative rates) of 95%.

While the correlation between observed and predicted values are extremely strong for both proxies, there were outliers in both cases. The identity of the outliers is mostly inconsistent between the two proxies, but *Papaver rhoeas* (the common poppy) is a highly problematic weed in cereal crops that had resistance evolvability underestimated by both models. *P. rhoeas* is the most common broadleaf weed in Europe (Torra et al 2010, 2018). It has, over the last 30 years, evolved resistance throughout Europe to a variety of herbicides with the ALS inhibitor (HRAC Group 2) and Auxin mimic (HRAC Group 4) modes of action, and it evolves resistance considerably more frequently and rapidly than its relatives within Ranunculales that are included within this dataset (*Fumaria densiflora* and *Ranunculus acris*). The consistent inability to predict the high degree of resistance evolvability in *P. rhoeas* may be due to the low number of closely related species in the IHRWD dataset and the long intervals of time separating these species (*Papaver* diverged from *Fumaria* 98.1 MYA and from *Ranunculus* 113 MYA (Li et al 2018). Such deep divergences allow greater variance of trait values to accumulate potentially leading to uncertainty in the predictions; in this case, *Papaver* has evolved a considerable increase in seed production and dormancy relative to the other Ranunculales in the dataset (Torra et al 2010; Rey-Caballero et al 2017), all of which increase its resistance evolvability. The under-estimation of resistance evolvability in *Vicia sativa* (which diverged from its nearest relative in the dataset 111.5 MYA) may also be attributed to a similar impact of deep divergence times between the tested members of larger clades.

Other taxa whose resistance evolvability is underestimated by the SVM models include *Amaranthus hybridus*, *Apera spica-venti*, *Bromus secalinus*, and *Digitaria insularis*. These species are members of larger genera and families that are well-represented in the datasets and are not separated from their near relatives in the dataset by deep divergence times. Therefore, alternative explanations for these species having greater resistance evolvability than may be predicted from their near relatives must be sought. It is not to be expected that these explanations are to be the same in each case; a great many traits have been suggested as affecting resistance evolvability. For example, an *EPSPS* triple mutation in *A. hybridus* has been suggested to give it greater levels of glyphosate resistance than other members of the genus (Perotti et al 2019; García et al 2019). It may also be related to them being more widespread weeds than their close relatives, or more common in agricultural settings, and therefore more likely to be treated with herbicides e.g. *Digitaria insularis* (sourgrass) is among the most competitive weed species in soybean and corn and is a frequent target of herbicides particularly in South America (de Carvalho et al. 2011; Lopez Ovejero et al 2017; Peterson et al 2018), as well as having a high dispersal ability to facilitate spread of resistance (Correia & Durigan 2009; de Carvalho et al. 2011; Lopez Ovejero et al 2017). Similarly, *Bromus secalinus* (Rye Brome) is particularly common across Europe and is problematic in winter crops (Davies et al 2020; Pytlarz & Gala-Czekaj 2022), so again is a more frequent target of herbicides.

The prediction methods validated here may be useful for identifying high-risk plants; as species become more problematic, the risk of resistance evolving rapidly and frequently may be useful for guiding weed management practices. The proxies employed, however, are very general, and therefore these analyses cannot predict resistance to specific herbicides or modes of action, or resistance in specific situations. Nevertheless, the principle validated here could be applied to such analyses, by including phylogeny along other predictors (e.g. Brocklehurst & Liu 2023) to identify species at greater risk under more precise circumstances.

### Resistance Regimes

With the demonstration of the power of phylogeny to predict a plant’s tendency to evolve resistance to herbicides, it follows that closely related taxa may be grouped into resistance regimes: regions of the phylogeny with similar patterns of resistance. Identifying such regimes can allow grouping of weeds into sets with similar resistance potential. Holt et al. (2013)’s assessment of plant families in which resistance is overrepresented can be thought of as an attempt to identify such regimes, albeit with *a priori* defined Linnaean taxonomic units. Phylogenetic comparative methods allow identification of such regimes without pre-defined locations on the phylogeny. The Surface algorithm is one such method, that locates a set of macroevolutionary regimes on a phylogeny, represented by evolution of a continuous trait under an Ornstein Uhlenbeck model with the trait being drawn to different optima in different regimes (Ingram & Mahler 2013). The practical value of the use of such regimes based on such broad proxies may be limited, and it would be more useful when combined with other predictors such as patterns within different herbicides and modes of action. The analysis here may be considered exploratory, and a demonstration of the potential value of more detailed work.

The broader pattern observed from these analyses is the separation of eudicots and monocots into distinct regimes with more frequent and rapid resistance evolution in the monocots (Figure 4). It implies the existence of fundamental evolutionary differences between these two clades that render one better able to evolve resistance than the other. There has been considerable discussion in the literature about why certain weeds evolve resistance more rapidly and more frequently than others, and differences between monocots and dicots could provide starting points for future analysis. Monocots appear to be particularly prone to metabolic (non-target site) resistance, potentially due to their more complex metabolic processes (Yu & Powles 2014; Nandula et al 2019). Large numbers of wind-dispersed seeds provide opportunities for resistance to spread, and have been cited as driving factors in large numbers of resistance cases in key genera (e.g. Tranel 2021; Prince & Carter 1985; Alcocer-Ruthling et al 1992; Warwick 1991; Chadha & Florentine 2021; Jhala et al 2021). Monocots have a significantly greater diversity of wind-pollinated taxa (Tiffney & Mazer 1995), providing greater opportunities for resistance genes to spread to new populations more rapidly. Monocots also have more relaxed constraints on their genome size and include species with larger genomes (Vinogradov 2001; Soltis et al 2003; Leitch et al. 2010). The precise relationship between increased genome size and evolutionary rate is unclear, but it has been suggested that larger genomes and duplicated genomes lead to greater rates of phenotypic evolution (Freeling & Thomas 2006; Van de Peer et al 2009; Kraaijeveld 2010).

Regime 4 provides an illustration of the distinction between the two proxies and the different aspects of “tendency to evolve resistance” that they are assessing. Species with large numbers of resistance cases will likely include those with a high potential to evolve resistance, but their abundance in specific countries and competitiveness in specific crop situations will determine how many different active ingredients they are exposed to and how frequently they are exposed to them. Therefore, the signal observed may be influenced to a certain extent by the species’ “weediness” as well as their intrinsic tendency to evolve resistance. The time-to-resistance proxy, on the other hand, assessed how rapidly resistance is likely to appear when a weed is exposed to a herbicide, regardless of how frequently this exposure takes place. Regime 4 includes helianth species such as *Xanthium strumarium* and *Helianthus annuus*. Both species have few cases of resistance (19 and 6 respectively; when subsampling is applied the resistance counts are 0.43 and 0.59 respectively), and although the majority of these cases were found in the USA, there are very few responses in the WSSA survey listing them as common or troublesome (Van Wychen 2019, 2020, 2021).

Nevertheless, their evolution of resistance is extremely rapid; the time to resistance in *H. annuus* is 7.5 and in *Xanthium strumarium* it is 4.5 years (the median across all species is 19 years). All helianth species included in this dataset are annuals, which could be a factor in their resistance evolvability due to short-lived species have a more rapid response to selection (Jasieniuk et al 1996, Holt et al 2013). *Ambrosia* is an invasive helianth weed that has become problematic in Europe and Asia (Zhou et al 2014; Oude Lansink et al 2018). Its success as an invasive species has in part been attributed to its rapid adaptive ability resulting from large amounts of genetic variability (Battlay et al 2023), which may also drive more rapid adaptation to herbicides. The patterns observed in this regime have implications for the treatment of weeds when they do become problematic: the fact that resistance is rare does not necessarily mean that resistance mitigation should not be a priority; on the rare occasions that these weeds do become problematic there is high potential for rapid resistance evolution.

## Conclusion

The complexity and variety of the factors that drive resistance makes predicting its evolution difficult, and there is no consensus over which drivers are most important. The recognition of phylogenetic signal in proxies for frequency and rapidity with which species evolve resistance indicates that a large driver of these resistance patterns are phenotypic traits of the weed species that have evolved at the clade level in deep time. Plant phylogeny therefore represents a tool for predicting which novel weed species are at greatest risk of resistance evolution, whether through machine learning approaches or identification of clades that represent resistance regimes. Phylogenetic comparative methods have uses in resistance research beyond those explored here, for example in more accurately assessing the correlation between resistance evolution and key “weedy” traits. With sufficient research, such methods may provide an invaluable addition to the numerous efforts to predict resistance evolution.

## Acknowledgements

We would like to thank Deepak Kaundun, Joe Downes, Deepmala Sehgal, Will Plumb, Melissa Brazier-Hicks, Breno Campos and Ian Heap for helpful comments and discussion, and Liam Revell and Graeme Lloyd for assistance with coding the analyses.

## Declarations

The authors declare no conflicts of interest

## Data Availability

All necessary data has been uploaded as electronic supplement

