## Supplementary material for "Phylogenetic signal in herbicide resistance evolvability, and the use of phylogenies to predict herbicide resistance risk": Figure S1

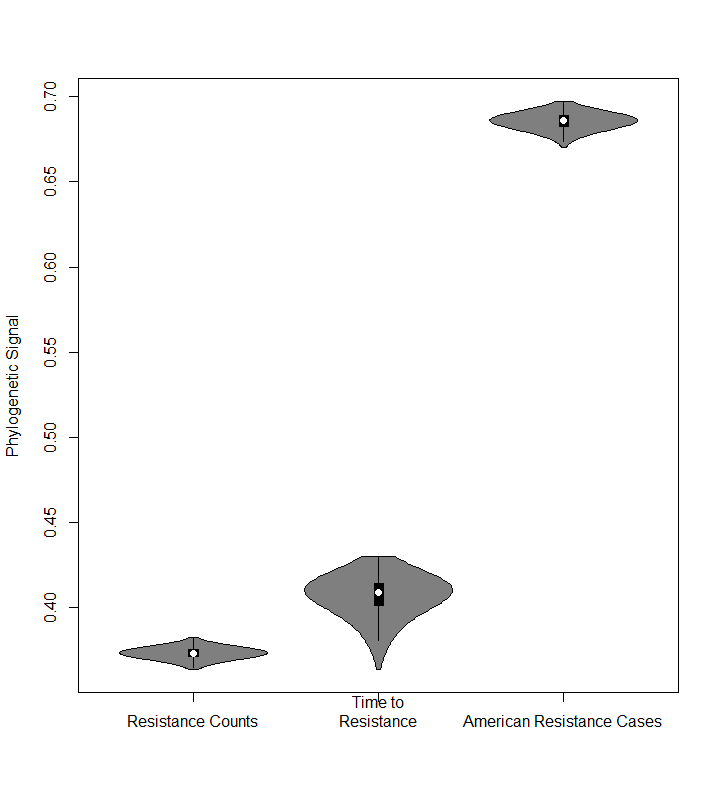

Fig S1: Phylogenetic signal of the resistance evolvability proxies, measured over 2000 trees randomly selected from the two birth death skyline analyses
